# Increasing Cas9-mediated homology-directed repair efficiency through covalent tethering of DNA repair template

**DOI:** 10.1101/231035

**Authors:** Eric J. Aird, Klaus N. Lovendahl, Amber St. Martin, Reuben S. Harris, Wendy R. Gordon

**Author notes:** Correspondence should be addressed to W.R.G.

## Abstract

The CRISPR-Cas9 system is a powerful genome-editing tool in which a guide RNA targets Cas9 to a site in the genome where the Cas9 nuclease then induces a double stranded break (DSB)^1,2^. The potential of CRISPR-Cas9 to deliver precise genome editing is hindered by the low efficiency of homology-directed repair (HDR), which is required to incorporate a donor DNA template encoding desired genome edits near the DSB^3,4^. We present a strategy to enhance HDR efficiency by covalently tethering a single-stranded donor oligonucleotide (ssODN) to the Cas9/guide RNA ribonucleoprotein (RNP) complex via a fused HUH endonuclease^5^, thus spatially and temporally co-localizing the DSB machinery and donor DNA. We demonstrate up to an 8-fold enhancement of HDR using several editing assays, including repair of a frameshift and in-frame insertions of protein tags. The improved HDR efficiency is observed in multiple cell types and target loci, and is more pronounced at low RNP concentrations.

## Introduction

Mammalian cells repair the DSB predominantly through two pathways: non-homologous end joining (NHEJ) or HDR^3^. The more frequent NHEJ pathway results in the formation of small insertions or deletions (indels) at the DSB site while the alternative HDR pathway can be utilized to insert exogenous DNA sequences into the genome^3,4^. For many applications, HDR is desired but is crippled by the low efficiency of recombination. Thus, various approaches have been developed to boost HDR frequency. Small molecules inhibiting NHEJ or upregulating HDR pathways have been reported to enhance HDR^6-8^. Another method to increase HDR efficiency is to regulate cell cycle progression or temporal control of Cas9 expression, either through small molecules or Cas9 protein fusions^9,10^. While these approaches and others have been reported to increase HDR efficiencies by as much as 10-fold, they suffer both from negative impacts on cell growth and inconsistencies between cell type and targeted loci^11^.

We reasoned that a strategy to covalently tether the DNA donor template to the RNP could dramatically enhance HDR efficiency by guaranteeing the presence of the donor DNA at the site of the DSB with the RNP complex, thus increasing the effective concentration of the donor template at the DSB and reducing the dimensionality of the reaction. Recent attempts to co-deliver the donor DNA using biotin-avidin^12^ and the SNAP tag^13^ suffer from additional steps required to modify the donor DNA. We recently showed that HUH endonucleases form robust covalent bonds with specific sequences of unmodified single-stranded DNA (ssDNA) and can function in fusion tags with diverse protein partners, including Cas9^5^. Formation of a phosphotyrosine bond between ssDNA and HUH endonucleases occurs within minutes at room temperature. Tethering the donor DNA template to Cas9 utilizing an HUH endonuclease could thus be a powerful approach to create a stable covalent RNP-ssODN complex.

## Results and discussion

We first fused the Porcine Circovirus 2 (PCV) Rep protein^5,14^ to either the amino (PCV-Cas9) or carboxyl (Cas9-PCV) terminus of Cas9 (**Fig. 1a**). We assayed for the selective attachment of ssDNA using fluorescently labelled ssDNA containing the recognition sequence for PCV (**Fig. 1b**). Both PCV fused Cas9 variants are able to bind the oligonucleotide while unfused Cas9 does not. In addition, mutating the catalytic tyrosine of PCV (Y96F) predictably prevents covalent attachment of the DNA. To assess reactivity of Cas9-PCV with ssODNs used in the HDR experiments, the upwards mobility shift of the protein-DNA conjugate was monitored using SDS-PAGE (**Fig. 1c**). An equimolar ratio of protein to DNA resulted in >60% covalent complex formation (by gel densitometry) after 15 minutes at room temperature. The ability of Cas9 to cleave DNA was assessed *in vitro* and found to not be perturbed by the addition of PCV at either terminus (**Supplementary Fig. 1**).

**Figure 1.**
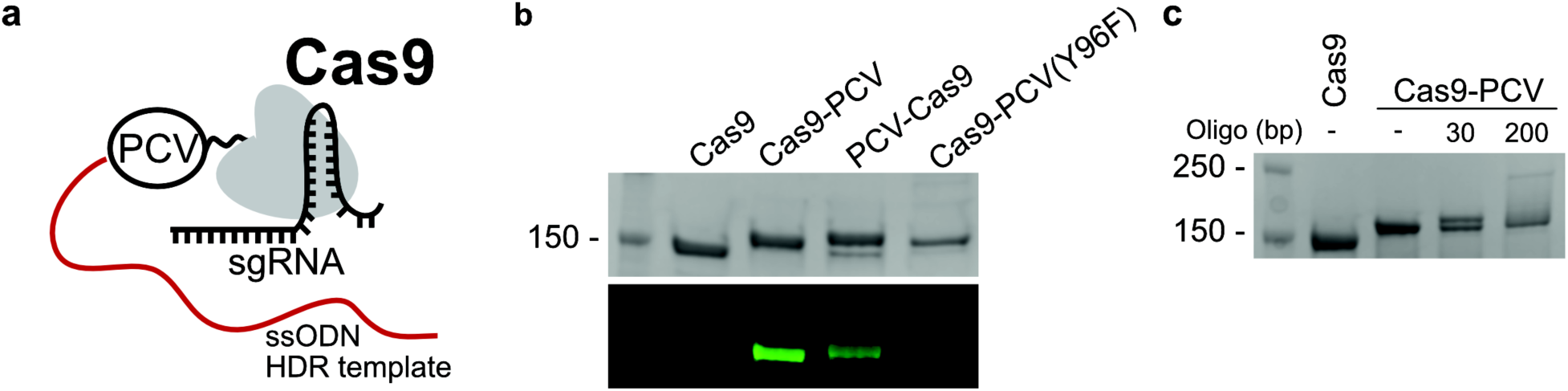
Selective covalent attachment of ssDNA to Cas9-PCV. (**a**) Schematic of Cas9 fused to the HUH endonuclease PCV with a covalently attached ssODN. (**b**) SDS-PAGE of Cas9 variants reacted with an Alexa 488 fluorescently labelled ssDNA containing the PCV recognition sequence. The top panel is the coomassie stained gel, and the bottom panel is the identical fluorescently imaged gel. PCV is fused to either the carboxyl (Cas9-PCV) or amino (PCV-Cas9) terminus of Cas9. Cas9-PCV(Y96F) represents catalytically inactive PCV(Y96F) fused to Cas9. (C) SDS-PAGE gel shift assay of Cas9 reacting with two ssDNA templates containing the PCV recognition sequence of differing lengths in a 1:1 ssDNA:Cas9 molar ratio.

We next used a recently described assay (Promega) to monitor the in-frame insertion of the 13 amino acid C-terminus of split-nanoluciferase (HiBiT) reporter into endogenous loci as a readout of HDR efficiency^15^. Adding the recombinant N-terminus (LgBiT) reconstitutes full-length nanoluciferase only in the edited cells and produces light upon addition of substrate. The intensity of light emitted is thus a relative measure of HDR efficiency. RNPs targeting the 3’ end of *GAPDH* were assembled *in vitro* and transfected into HEK-293T cells with or without ssODN containing the PCV recognition sequence (**Fig. 2a**). When using an ssODN lacking the PCV recognition sequence (PCV-ssODN), all versions of Cas9 resulted in similar luminescence levels when assayed 48 hours post-transfection (**Fig. 2b**). However, upon addition of an ssODN containing the PCV recognition sequence (PCV+ ssODN), a significant two to three-fold change in luminescence is observed for cells transfected with Cas9-PCV fusions (**Fig. 2c**). The increase in luminescence was abrogated in catalytically inactive PCV (Y96F) fused to Cas9, suggesting the specific attachment of ssODN to Cas9 via PCV is responsible for the increase in HDR frequency. This enhancement of HDR is not limited to HEK-293T cells or the particular *GAPDH* locus. We have targeted *GAPDH* in osteosarcoma U2-OS cells for HiBiT insertion and observed similar effects (**Fig. 2d**).We have also targeted an in-frame locus between two domains of vinculin, which is expressed at lower levels than GAPDH in HEK293T cells, and measured a significant increase in luminescence using the Cas9-PCV fusion in comparison to Cas9 alone (**Fig. 2e**). Additionally, combining our methodology with the addition of SCR7, an NHEJ inhibitor, increases HDR efficiency further (**Supplementary Fig. 2**), suggesting that covalent attachment of DNA could be combined with other methods to create additive effects.

**Figure 2.**
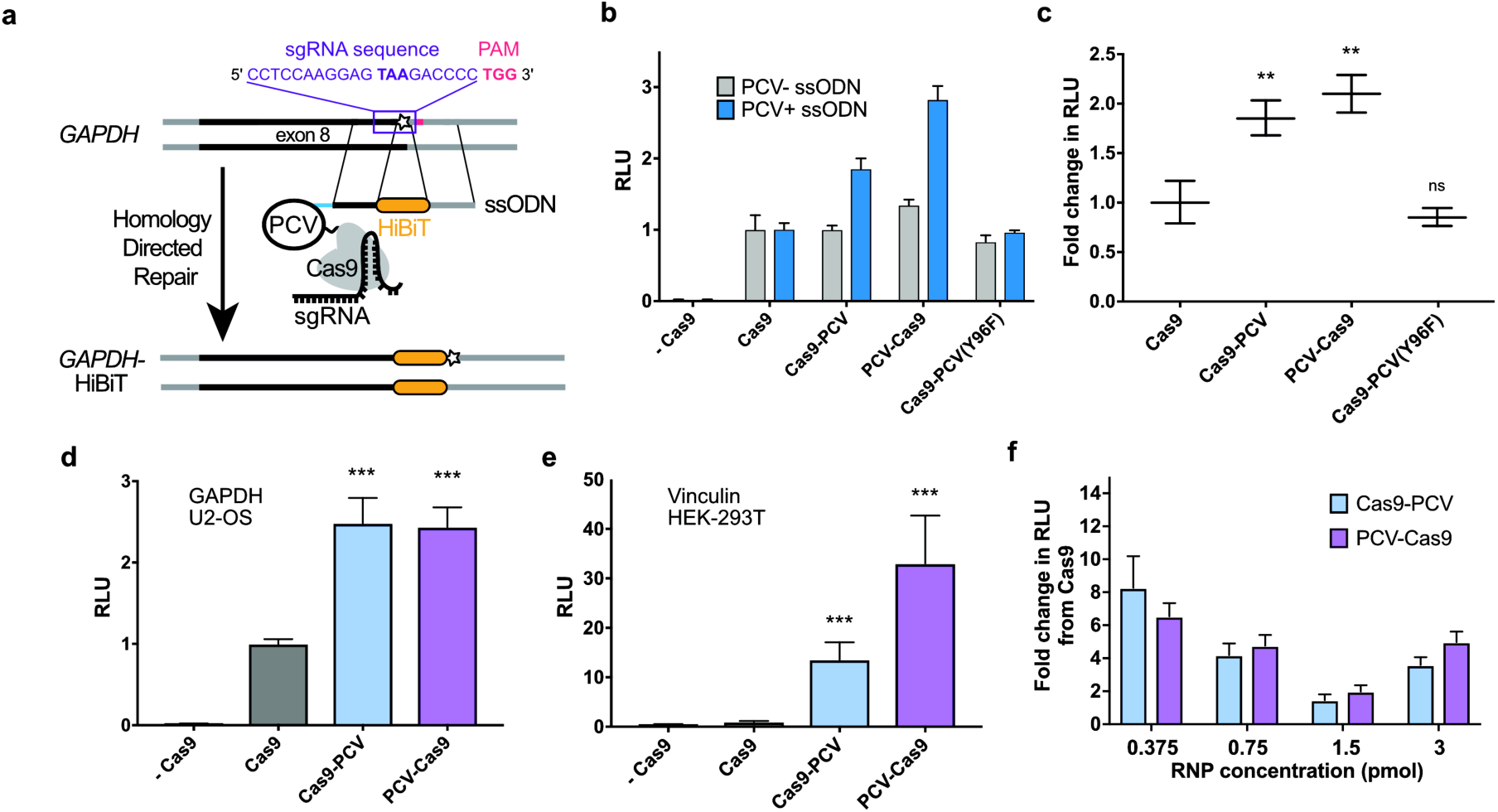
Covalent attachment of ssODN to Cas9-PCV enhances HDR. (**a**) Schematic of split luciferase insertion. The C-terminus of NanoLuc nanoluciferase (HiBiT) is encoded on the 200bp ssODN along with the 5’ PCV recognition sequence and targeted to the 3’ end of *GAPDH*. (**b**) Assaying luminescence using different Cas9 variants when inserting HiBiT into *GAPDH* in HEK-293T cells. PCV is fused to either the amino (PCV-Cas9) or carboxyl (Cas9-PCV) terminus of Cas9. Transfections were performed with ssODN lacking the PCV recognition sequence (PCV- ssODN) or ssODN containing the PCV recognition sequence (PCV+ ssODN). Units are displayed in relative light units (RLU) normalized to Cas9. (**c**) The calculated fold change from (b) between the PCV-ssODN and PCV+ ssODN is shown for each variant. (**d**) Targeting the *GAPDH* locus in U2-OS cells. (**e**) Targeting a locus in vinculin in HEK-293T cells using an ssODN containing the PCV recognition sequence. (**f**) Fold change in RLU compared to Cas9 when varying the amount of RNP (equimolar ssODN). All graphs represent data from one of multiple independent experiments exhibiting similar results. Data are shown as mean +/− SD (n=3). Significance calculated using 2-tailed Student’s t-test: ** P < 0.01, *** P < 0.001, ns = no significance (P >0.05).

Importantly, the HDR enhancement observed in the context of PCV-tagged Cas9 and co-delivery of the repair DNA cannot be recapitulated by merely increasing the Cas9 or donor DNA concentrations. RNP titrations reveal increased HDR for Cas9-PCV when compared to Cas9 regardless of the concentration (**Fig. 2f**). Interestingly, the enhancement is much more pronounced at lower concentrations of Cas9-PCV, where up to an 8-fold change in HDR exists. Similarly, increasing the concentration of ssODN containing the PCV recognition sequence has no effect on Cas9 HDR efficiency while for Cas9-PCV, the maximal luminescence readout is seen when at a 1 to 1 ratio of RNP to ssODN (**Supplementary Fig. 2**). Taken together, the luminescence data strongly suggests the significant increase in HDR is due to covalent tethering of the DNA template to Cas9-PCV.

We used quantitative-PCR (qPCR) to confirm that the effect seen at the protein level in the luminescence assay is consistent at the DNA level. We assayed genomic DNA preparations for insertion of HiBiT. Primers were designed to amplify only in the presence of the insertion with a reference primer pair located upstream on *GAPDH* (**Supplementary Fig. 3**). The upstream primer for the HiBiT pair binds outside of the ssODN sequence so no amplification occurs from the ssODN itself. Using the Pfaffl method to quantitate differences in cycle threshold values^16^, we observed a consistent 2-fold change in HDR efficiency between Cas9 and Cas9-PCV fusions that mirrors the luminescence results (**Supplementary Fig. 3**).

Next, we aimed to quantify HDR using an assay providing absolute instead of relative quantitation of HDR while also performing a different type of repair. Using an integrated dual fluorescent GFP-mCherry reporter HEK-293T cell line^17^, we restored mCherry fluorescence by providing an ssODN HDR template to correct a frameshift mutation (**Fig. 3a and b**). Using flow cytometry, the HDR efficiency measured for cells transfected with Cas9-PCV fusions and PCV+ ssODN was found to be significantly higher than unfused Cas9 at 3 pmol RNP/ssODN (**Fig. 3c**). When decreasing the covalent complex concentration, HDR enhancement is 6-8 fold while Cas9-PCV mediated editing efficiency remains relatively constant. These findings are consistent with the luminescence assay data where lower concentrations result in increased enhancement. We also transfected RNPs without ssODN to account for indel formations that can also produce an in-frame gene. The degree of mCherry restoration without ssODN is relatively constant amongst the Cas9 variants while the contribution when adding ssODN for Cas9-PCV fusions is significantly greater than for Cas9 (**Fig. 3d**).

**Figure 3.**
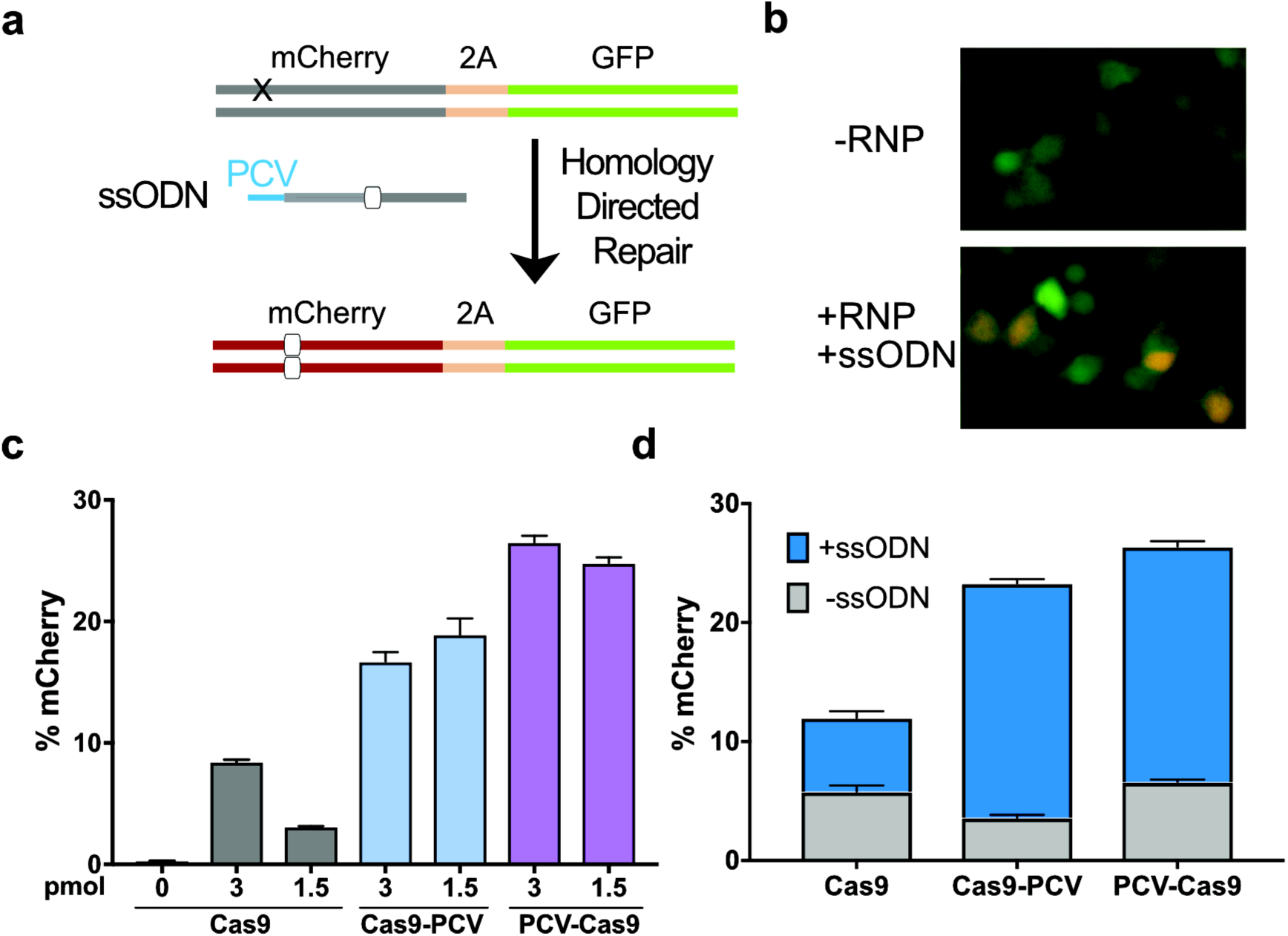
Covalent complex of ssODN and RNP enhances HDR efficiency 6-8 fold in a fluorescent reporter cell line. (**a**) HEK-293T cells stably expressing a mutant mCherry-GFP reporter are edited by HDR through a frameshift correction, restoring mCherry activity. (**b**) Representative microscopy images of fluorescent reporter editing. (**c**) The percent of mCherry positive cells determined by flow cytometry at two different RNP concentrations using an ssODN containing the PCV recognition sequence (**d**) RNP transfections at 3 pmol in the presence or absence of ssODN. Data are shown as mean +/− SD (n=3). For (c) and (d), the statistical significance of %mCherry positive cells between PCV-fusions of Cas9 and Cas9 alone was <0.001, calculated using 2-tailed Student’s t-test.

We have presented a strategy to improve HDR that can easily be incorporated into any Cas9 workflow utilizing RNPs. By simply appending a 13 basepair recognition sequence to the 5’ end of an ssODN and using a Cas9-PCV fusion protein, HDR efficiency can be increased up to 8 fold as measured by three separate approaches.Importantly, significantly lower concentrations of HUH-tagged Cas9 are needed to achieve high efficiencies of HDR, which may offer advantages for downstream clinical uses. Collectively, this methodology of covalently tethering the donor DNA template to Cas9 is broadly applicable and effective at enhancing HDR across various cell types and targeting loci, and may be further improved by varying PCV fusion location and optimizing homology arm lengths^18,19^. The ability to co-localize the donor DNA may offer some advantages in contexts where RNPs are delivered via injection and the strategy should also be applicable to double stranded templates with short single-stranded overhangs. Moreover, the use of additional HUH-tags with different target sequences may allow multiplexed one-pot reactions due to the orthogonality of HUH endonucleases. Our method is also easily scalable for RNP-based *in vivo* applications as no chemical modifications are needed on the DNA template. Furthermore, this technique does not require modulating activity of any endogenous protein as is common with small molecule HDR enhancers. We have also shown that different approaches to enhancing HDR can be combined with DNA tethering to further enhance editing efficiency.

## Acknowledgements

We would like to thank Marie Schwinn and Keith Wood at Promega for providing the split-luciferase insertion protocol, sequences, and reagents prior to commercial release. We would like to thank Eric Hendrickson, Brian Ruis, and David Largaespada for helpful discussions and Amanda Hayward for critical reading of the manuscript. We would like to thank Hideki Aihara for use of his plate reader and qPCR machine and the University Imaging Centers at the University of Minnesota for support.

## Author contributions

E.J.A. and W.R.G. designed experiments, E.J.A. and A.S. performed experiments and analysis, E.J.A. and K.N.L designed and constructed fusion constructs, A.S. and R.S.H designed and created reporter cell line, and E.J.A, K.N.L, and W.R.G assembled the manuscript with contributions from all authors.

## Methods

### Nucleotide sequences

The sgRNA, ssODN, and primer sequences are found in **Supplementary Table 1**. In ssODNs lacking the PCV recognition sequence, an equal length (13 bp) randomized sequence was added in place so homology arm length was kept consistent. All ssODNs and primers synthesized by Integrated DNA Technologies.

### Construct design

*Streptococcus pyogenes* Cas9 was amplified out of the plasmid pET15_SP-Cas9 (a gift from Niels Geijsen, Addgene plasmid #62731) and inserted in pTD68_SUMO-PCV2 at the BamHI site using Infusion cloning (Clontech) to create C-terminally fused Cas9-PCV. N-terminally fused PCV-Cas9 was constructed using golden gate assembly in pTD68_SUMO with insertion of an H4-2 linker between PCV and Cas9^20^. Protein sequences can be found in Supplementary Table 1. Catalytically dead Cas9-PCV (Y96F) was created by QuikChange II site directed mutagenesis kit (Agilent Technologies).

### Protein expression and purification

Bacterial expression of proteins was performed in BL21(DE3) *E. coli* using an autoinduction protocol^21^. Following inoculation, cells were rotated at 300 rpm for 8 hours at 37°C, followed by 24 hours at 24°C. Protein purification was performed using the methods of Anders and Jinek^22^. Cells were pelleted and lysed in 50 mM Tris, 200 mM NaCl, 20% sucrose, pH 7.4. The clarified supernatant was first passed over Ni-NTA resin (Thermo Scientific), washed with 15 column volumes of 50 mM Tris, 200 mM NaCl, 15 mM imidazole, pH 7.4, and eluted with 250 mM imidazole followed by overnight incubation with SUMO protease Ulp1 at 4°C while dialyzing into 50 mM HEPES, 150 mM KCl, 10% glycerol, 1 mM DTT, 1 mM EDTA, pH 7.5. Cation exchange chromatography was then performed using a 1 mL HiTrap SP HP column (GE Healthcare), eluting using a gradient from 100 mM to 1 M KCl. Fractions were analyzed by SDS-PAGE and pooled. Protein was concentrated in a 100 kDa MWCO spin concentrator (Amicon). Final protein concentration was measured by Bradford Assay (Bio-Rad) or A_280_ absorbance. Aliquots were stored at −20°C or flash frozen in dry ice/IPA.

### Covalent DNA attachment to Cas9-PCV

Equimolar amounts of Cas9-PCV and the sequence specific ssODN were incubated at room temperature for 10-15 minutes in Opti-MEM (Corning) supplemented with 1 mM MgCl_2_. Confirmation of the linkage was analyzed by SDS-PAGE. For the fluorescent oligonucleotide reactions, 1.5 pmol of Alexa 488-conjugated ssDNA (IDT) was incubated with 1.5 pmol Cas9-PCV in the above conditions and separated by SDS-PAGE. Gels were imaged using a 473 nm laser excitation on a Typhoon FLA9500 (GE) at the University of Minnesota Imaging Center.

### *In vitro* cleavage assay

An eGFP-pcDNA3 vector was linearized with BsaI (NEB). 30 nM sgRNA targeting GFP, 30nM Cas9, and 1× T4 ligase buffer were incubated for 10 minutes prior to adding linearized DNA to a final concentration of 3 nM. The reaction was incubated at 37°C for 1 to 24 hours, then separated by agarose gel electrophoresis and imaged using SYBR safe gel stain (Thermo Fisher). The percent cleaved was calculated by comparing densities of the uncleaved band and the top cleaved band using Image Lab software (Bio-Rad).

### Cell culture

HEK-293T, U2-OS, and stable dual-fluorescent HEK-293T cells were cultured in DMEM (Corning) supplemented with 10% FBS (Gibco) and 0.5% penicillin/streptomycin (Gibco). Cells were incubated at 37°C in 5% CO_2_.

### RNP transfection

Guide RNA sequences were purchased from Integrated DNA Technologies. Guide RNA was formed by mixing equimolar amounts of tracrRNA and crisprRNA in duplex buffer (IDT), heating to 95°C for 5 minutes, and cooled on the benchtop. Equimolar amounts of Cas9 and guide RNA were incubated in Opti-MEM supplemented with 1 mM MgCl_2_ for 5-10 minutes at room temperature. Following RNP formation, equimolar amount of ssODN was added and incubated for 10-15 minutes at room temperature. Transfections were carried out with RNAiMAX (Invitrogen) using the manufacturer’s suggested protocol in either 96 well plate format with 1.5 pmol RNP/ssODN and 4×10^4^ viable cells per well or 24 well plate using 3 pmol RNP/ssODN and 2×10^5^ viable cells per well unless noted. For SCR7-containing experiments, 1 μM SCR7 (Sigma) was added at the time of transfection. Cells were washed and fresh media was added 24 hours later.

### Luminescence activity assay

24-48 hours post-transfection, luminescence measurements were performed using the Nano-Glo HiBiT lytic assay system (Promega). Briefly, confluent cells were detached and lysed in a buffer containing the recombinant N-terminus of nanoluciferase (LgBiT) and nanoluciferase substrate furimazine in 96-well half-volume plates. Lysates were incubated for 10 minutes rotating, and luminescence intensities were recorded on a LMax II luminometer (Molecular Devices) at a one second integration time.

### Quantitative PCR

48 hours post-transfection, cells from individual 96 plate wells were spun down at 500x g for 5 minutes. Genomic DNA was extracted and purified from cell pellets using Purelink genomic DNA purification kit (Invitrogen). In triplicate, SYBR-based qPCR reactions were performed using PowerUp SYBR Green Master Mix (Applied Biosciences) on a Bio-Rad CFX96 real-time thermocycler. Cycle conditions: 95°C for 2 minutes; 35 cycles of 95°C for 15 seconds and 58°C for 45 seconds. Melt curve step at 0.5°C per minute. Data was analyzed on CFX Manager software (Bio-Rad). Relative changes in abundance between HiBiT containing DNA and the GAPDH reference were calculated using the Pfaffl method^16^. Melt curve analysis and primer amplification efficiency measurements on serial diluted DNA templates were also carried out (**Supplementary Fig 3**).

### mCherry-GFP HEK-293T editing

Transfections carried out as described above in 24 well plates with 3 pmol of RNP/ssODN unless stated. 48 hours post-transfection, cells were detached and centrifuged at 500× g for 5 minutes. Following aspiration of supernatant, cells were resuspended in 200 μl PBS + 1% EDTA and transferred to a 96 well plate. 10,000 to 100,000 cells were analyzed using flow cytometry on a BD Biosciences FACSCalibur with data collected using BD FACSDiva and compiled using FlowJo (version 10.4). Cells were initially gated based on FSC/SSC and then gated on presence or lack of fluorescent protein expression (**Supplementary Fig. 4**). In separate experiments, fluorescence microscopy cell images were obtained 24 hours post-transfection on an EVOS FL Auto (Life Technologies) using a 20× objective. GFP and RFP light cubes were used to image GFP and mCherry, respectively. Images were processed using Fiji (version 1.51 r).

### Data availability

Additional supporting data beyond what is in the text and supplementary information is available upon request.

